# *Vcfanno*: fast, flexible annotation of genetic variants

**DOI:** 10.1101/041863

**Authors:** Brent S. Pedersen, Ryan M. Layer, Aaron R. Quinlan

**Affiliations:** Department of Human Genetics, University of Utah, Salt Lake City, UT 84105; USTAR Center for Genetic Discovery, University of Utah, Salt Lake City, UT 84105; Department of Biomedical Informatics, University of Utah, Salt Lake City, UT 84105

**Keywords:** genetic variation, SNP, annotation, VCF, variant, variant prioritization, genome analysis

## Abstract

**Background:** The integration of genome annotations and reference databases is critical to the identification of genetic variants that may be of interest in studies of disease or other traits. However, comprehensive variant annotation with diverse file formats is difficult with existing methods.

**Results:** We have developed *vcfanno* as a flexible toolset that simplifies the annotation of genetic variants in VCF format. *Vcfanno* can extract and summarize multiple attributes from one or more annotation files and append the resulting annotations to the INFO field of the original VCF file. *Vcfanno* also integrates the lua scripting language so that users can easily develop custom annotations and metrics. By leveraging a new parallel “chromosome sweeping” algorithm, it enables rapid annotation of both whole-exome and whole-genome datasets. We demonstrate this performance by annotating over 85.3 million variants in less than 17 minutes (>85,000 variants per second) with 50 attributes from 17 commonly used genome annotation resources.

**Conclusions:** *Vcfanno* is a flexible software package that provides researchers with the ability to annotate genetic variation with a wide range of datasets and reference databases in diverse genomic formats.

**Availability:** The *vcfanno* source code is available at https://github.com/brentp/vcfanno under the MIT license, and platform-specific binaries are available at https://github.com/brentp/vcfanno/releases. Detailed documentation is available at http://brentp.github.io/vcfanno/, and the code underlying the analyses presented can be found at https://github.com/brentp/vcfanno/tree/master/scripts/paper.

## BACKGROUND

The VCF files[1] produced by software such as GATK[2] and FreeBayes[3] report the polymorphic loci observed among a cohort of individuals. Aside from the chromosomal location and observed alleles, these loci are essentially anonymous. Until they are embellished with genome annotations, it is nearly impossible to quickly answer basic questions such as “was this variant seen in ClinVar,” or “what is the alternate allele frequency observed in the 1000 Genomes Project?” There is an extensive and growing number of publicly available annotation resources (Ensembl, UCSC) and reference databases of genetic variation (e.g., ClinVar, Exome Aggregation Consortium (ExAC), 1000 Genomes) that provide context that is crucial to variant interpretation. It is also common for individual labs and research consortia to curate custom databases that are used, for example, to exclude variants arising in genes or exons which are systematic sources of false positives in exome or genome resequencing studies. Other annotations, such as low-complexity regions[4], transcription factor binding sites, regulatory regions, or replication timing[5] can further inform the prioritization of genetic variants related to a phenotype. The integration of such annotations is complementary to the gene-based approaches provided by snpEff[6], Annovar[7], and VEP[8]. Each of these tools can provide additional, region-based annotation, yet they are limited to the genome annotation sets provided by the software. While extensive variant annotation is fundamental to nearly every modern study of genetic variation, no existing software can flexibly and simply annotate VCF files with so many diverse data sets.

We have therefore developed *vcfanno* as a fast and general solution for variant annotation that allows variants to be “decorated” with any annotation dataset in common genomics formats. In addition to providing the first method that is capable of annotating with multiple annotation sets at a time, *vcfanno* also avoids common issues such as inconsistent chromosome labeling (“chr1” vs. “1”) and ordering (*1,2,…10…*, or *1,10,11…*) among the VCF and annotation files. To maximize performance with dozens of annotation files comprised of millions of genome intervals, we introduce a parallel sweeping algorithm with high scalability. In an effort to make *vcfanno*’s annotation functionality as flexible as possible, we have also embedded a lua (www.lua.org) scripting engine that allows users to write custom operations.

## METHODS

### Overview of the *vcfanno* functionality

*Vcfanno* annotates variants in a VCF file (the “query” intervals) with information aggregated from the set of intersecting intervals among many different annotation files (the “database” intervals) stored in common genomic formats such as BED, GFF, GTF, VCF, and BAM. It utilizes a “streaming” intersection algorithm that leverages sorted input files to greatly reduce memory consumption and improve speed. As the streaming intersection is performed (details below), database intervals are associated with a query interval if there is an interval intersection. Once all intersections for a particular query interval are known, the annotation proceeds according to user-defined operations that are applied to the attributes (e.g., the “score” column in a BED annotation file or an attribute in the INFO field of a VCF annotation file) data within the database intervals. As a simple example, consider a query VCF of single nucleotide variants (SNVs) that was annotated by SNVs from an annotation database such as a VCF file of the dbSNP resource. In this case, the query and database variants are matched on position, REF, and ALT fields when available, and a value from the overlapping database interval (e.g., minor allele frequency) is carried forward to become the annotation stored in the INFO field of the query VCF. In a more complex scenario where a query structural variant intersects multiple annotation intervals from each database, the information from those intervals must be aggregated. One may wish to report each of the attributes as a comma-separated list via the ‘concat’ operation. Alternatively, one could select the maximum allele frequency via the ‘max’ operation. For cases where only a single database interval is associated with the query, the choice of operation will not affect the summarized value.

An example VCF INFO field from a single variant before and after annotation with *vcfanno* is shown in **Figure 1**. A simple configuration file is used to specify both the source files and the set of attributes (in the case of VCF) or columns (in the case of BED or other tab-delimited) that should be added to the query file. In addition, the configuration file allows annotations to be renamed in the resulting VCF INFO field. For example, we can extract the allele frequency (AF) attribute from the ExAC VCF file[9] and rename it as “exac_aaf” in the INFO field of the VCF query records. The configuration file allows one to extract as many attributes as needed from any number of annotation datasets.

**Figure 1.**
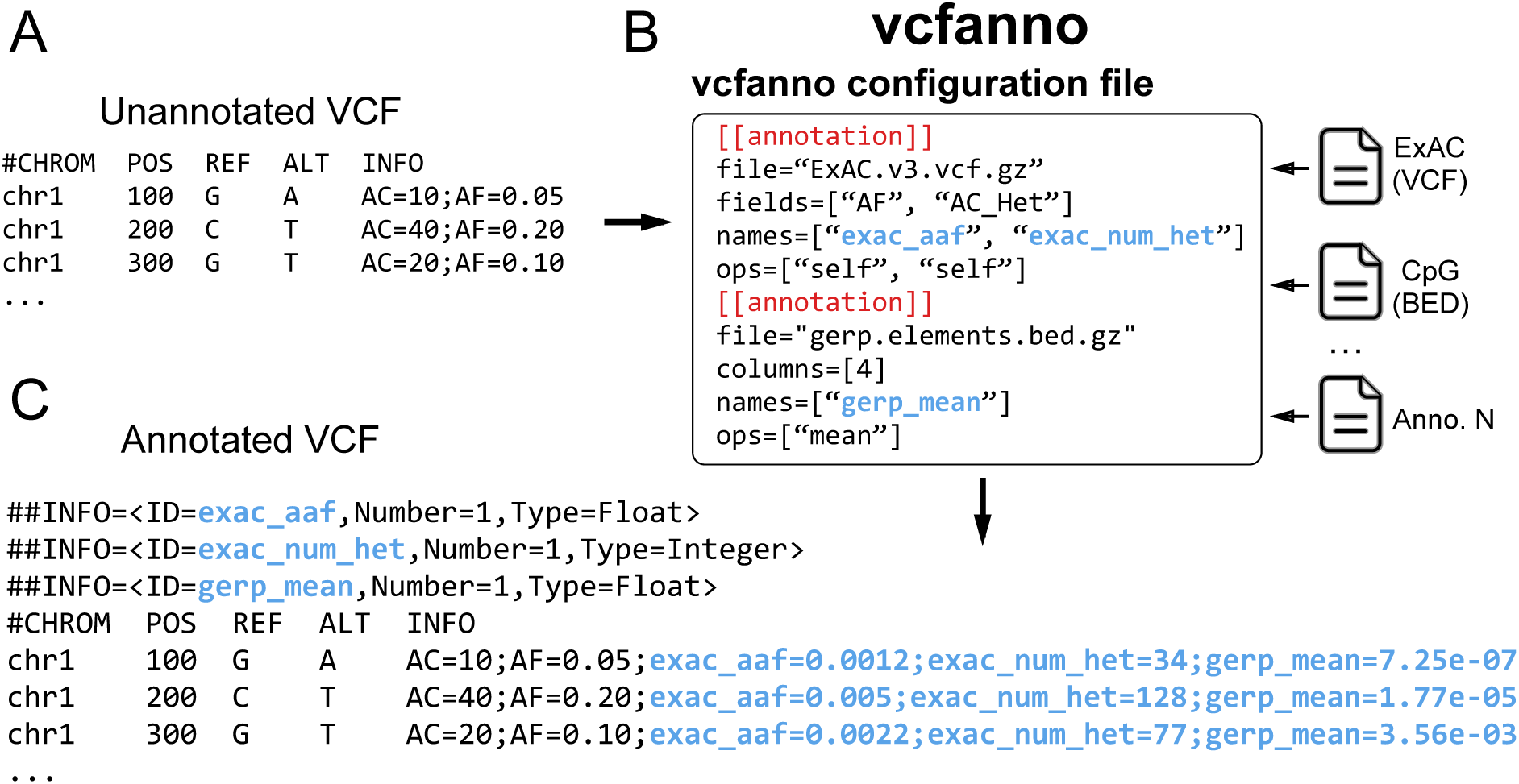
Overview of the *vcfanno* workflow. An unannotated VCF (A) is sent to *vcfanno* (B) along with a configuration file that indicates the paths to the annotation files, the attributes to extract from each file, and the methods that should be used to describe or summarize the values pulled from those files. The new annotations in the resulting VCF (C) are shown in blue text with additional fields added to the INFO column.

### Overview of the chrom-sweep algorithm

The chromosome sweeping algorithm (“chrom-sweep”) is an adaptation of the streaming, sort-merge join algorithm, and is capable of efficiently detecting interval intersections among multiple interval files, as long as they are sorted by both chromosome and interval start position. Utilized by both BEDTOOLS[10, 11] and BEDOPS[12], chrom-sweep finds intersections in a single pass by advancing pointers in each file that are synchronized by genomic position. At each step in the sweep, these pointers maintain the set of intervals that intersect a particular position and, in turn, intersect each other. This strategy is advantageous for large datasets because it avoids the use of data structures such as interval trees or hierarchical bins (e.g., the UCSC binning algorithm[13]). While these tree and binning techniques do not require sorted input, the memory footprint of these methods scales poorly, especially when compared to streaming algorithms, which typically exhibit low, average-case memory needs.

The chrom-sweep algorithm implemented in *vcfanno* proceeds as follows. First, we create an iterator of interval records for the query VCF and for each database annotation file. We then merge intervals from the query VCF and each annotation into a single priority queue, which orders the intervals from all files by chromosome and start coordinate, while also tracking the file from which each interval came. *Vcfanno* progresses by requesting an interval from the priority queue and inserts it into a cache. If the most recently observed interval is from the query VCF, we check for intersections with all database intervals that are currently in the cache. Since *vcfanno* requires that all files be sorted, we know that intervals are entering the cache ordered by start coordinate. Therefore, in order to check for overlap, we only need to check that the start of the new interval is less than the end of any of the intervals in the cache (assuming half-open intervals). An example of the sweeping algorithm is shown in **Figure 2** for a case involving 2 annotation files and 3 records from a single query VCF. The contents of the cache are shown as the sweep reaches the start of each new interval. When a new query interval enters the cache, any interval that does not intersect it is ejected from the cache. If the removed interval originated from the query VCF, it is sent, together with each of the intersecting annotation intervals to be processed according to the operations specified in the configuration file. The resulting annotations are stored in the INFO field of the VCF file and the updated VCF record is reported as output.

**Figure 2.**
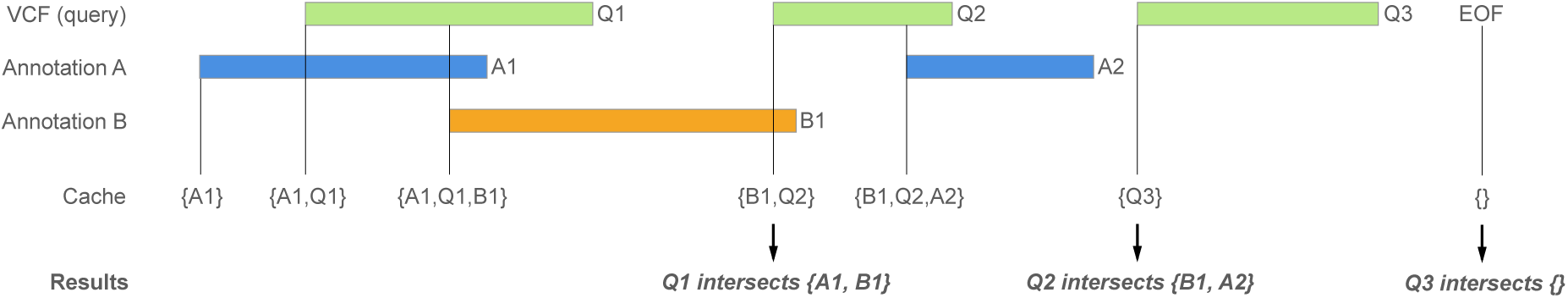
Overview of the chrom-sweep interval intersection algorithm. The chrom-sweep algorithm sweeps from left to right as it progresses along each chromosome. Green intervals from the query VCF in the 1st row are annotated by annotation files A (blue) and B (orange) in the 2nd and 3rd rows, respectively. The cache row indicates which intervals are currently in the cache at each point in the progression of the sweeping algorithm. Intervals enter the cache in order of their chromosomal start position. First A1 enters the cache followed by Q1. Since Q1 intersects A1, they are associated; as are Q1 and B1 when B1 enters the cache. Each time a new query interval enters the cache, any interval it does not intersect is ejected. Therefore, when Q2 enters the cache, Q1 and A1 are ejected. Since Q1 is a query interval, it is sent to be reported as output. Proceeding to the right, A2 and then Q3 enter the cache; the latter is a query interval and so the intervals that do not overlap it – B1, Q2, and A2 – are ejected from the cache with the query interval, Q2, which is sent to the caller. Finally, as we reach the end of the incoming intervals, we clear out the final Q3 interval and finalize the output for this chromosome.

### Limitations of the chrom-sweep algorithm

Owing to the fact that annotation sets are not loaded into memory intensive data structures, the chrom-sweep algorithm easily scales to large datasets. However, it does have some important limitations. First, it requires that all intervals from all annotation files adhere to the same chromosome order. While conceptually simple, this is especially onerous since VCFs produced by variant callers such as GATK impose a different chromosome order (1, 2,…21, X, Y, MT) than most other numerically sorted annotation files, which would put MT before X and Y. Of course, sorting the numeric chromosomes as characters or integers also results in different sort orders. Discrepancies in chromosome ordering among files are often not detected until substantial computation has already been performed. A related problem is when one file contains intervals from a given chromosome that the other does not, it’s not possible to distinguish whether the chromosome order is different or if that chromosome is simply not present in one of the files until all intervals are parsed.

Second, the standard chrom-sweep implementation is sub-optimal because it is often forced to consider (and parse) many annotation intervals that will never intersect the query intervals, resulting in unnecessary work[14]. For example, given a VCF file of variants that are sparsely distributed throughout the genome (e.g., a VCF from a single exome study) and dense data sets of whole-genome annotations, chrom-sweep must parse and test each interval of the whole-genome annotations for intersection with a query interval, even though the areas of interest comprise less than 1% of the regions in the file. In other words, sparse queries with dense annotation files represent a worst-case scenario for the performance of chrom-sweep because a high proportion of the intervals in the data sets will never intersect.

A third limitation of the chrom-sweep algorithm is that, due to the inherently serial nature of the algorithm, it is difficult to parallelize the detection of interval intersections and the single CPU performance is limited by the speed at which intervals can be parsed. Since the intervals arrive in sorted order, skipping ahead to process a new region from each file in a different processing thread is difficult without a pre-computed spatial index of the intervals, and reporting the intervals in sorted order after intersection requires additional bookkeeping.

### A parallel chrom-sweep algorithm

To address these shortcomings, we developed a parallel algorithm that concurrently chrom-sweeps “chunks” of query and database intervals. Unlike previous in-memory parallel sweeping methods that uniformly partition the input[15], we define (without the need for preprocessing[16]) chunks by consecutive query intervals that meet one of two criteria: either the set reaches the “chunk size” threshold, or the genomic distance to the next interval exceeds the “gap size” threshold. Restricting the chunk size creates reasonably even work among the threads to support efficient load balancing (i.e., to avoid task divergence). The gap size cutoff is designed to avoid processing an excessive number of unrelated database intervals that reside between distant query intervals.

As soon as a chunk is defined, it is scheduled to be swept in parallel along with the other previously defined chunks. The bounds of the query intervals in the chunk determine the bounds of the intervals requested from each annotation file (**Figure 3**). Currently these requests are to either a Tabix indexed file or a BAM file via the bíogo package[17], but any spatial query can be easily supported. An important side effect of gathering database intervals using these requests is that, while the annotations files must be sorted, there is no need for the chromosome orders of the annotations to match. This, along with internally removing any “chr” prefix, alleviates the associated chromosome order and representation complexities detailed above. Conceptually, the set of intervals from these requests are combined with the query intervals to complete the chunk, which is then processed by the standard chrom-sweep algorithm. However, in practice this is accomplished by streams so that only the query intervals are held in memory while the annotation intervals are retrieved from their iterators during the chrom-sweep. One performance bottleneck in this strategy is that the output should be sorted, and since chunks may finish in any order, we must buffer completed chunks to restore sorted order. This, along with disk speed limitations, is the primary source of overhead preventing optimal parallelization efficiency.

**Figure 3.**
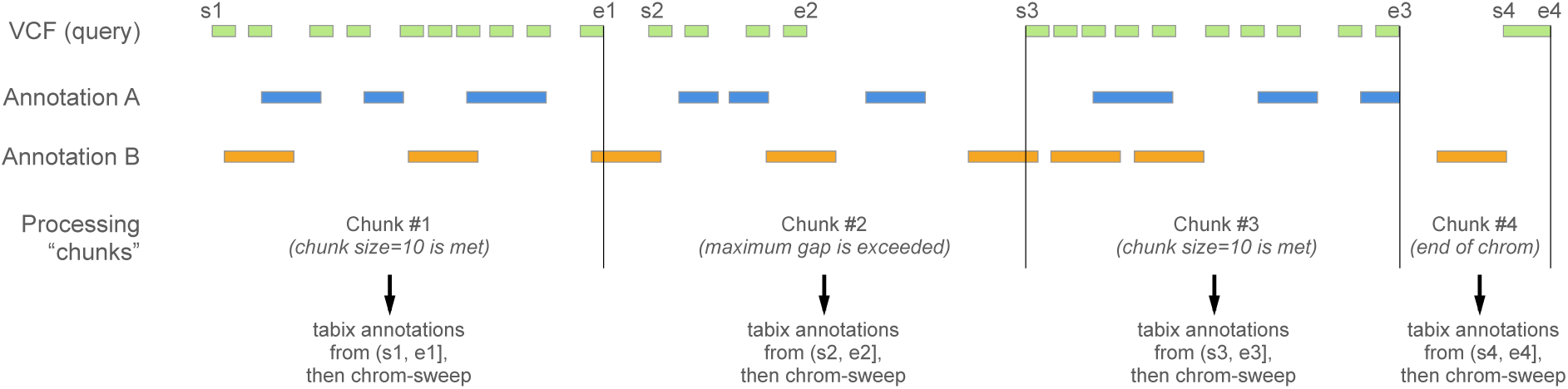
Parallel sweeping algorithm. As in Figure 2, we sweep across the chromosome from lower to higher positions (and left to right in the figure). The green query intervals are to be annotated with the 2 annotation files depicted with blue and orange intervals. The parallelization occurs in *chunks* of query intervals delineated by the black vertical lines. One process reads query intervals into memory until a maximum gap size to the next interval is reached (e.g. chunk 2, 4), the number of intervals exceeds the chunk size threshold (e.g. chunks 1, 3). While a new set of query intervals accumulates, the first chunk, bounded to the right by the first vertical black line above, is sent for sweeping and a placeholder is put into a FIFO (first-in, first-out) queue, so that the output remains sorted even though other chunks may finish first. The annotation files are queried with regions based on the bounds of intervals in the query chunk. The queries then return streams of intervals, and finally those streams are sent to the chrom-sweep algorithm in a new process. When it finishes, its placeholder can be pulled from the FIFO queue and the results are yielded for output.

### *Vcfanno* implementation

*Vcfanno* is written in Go (https://golang.org), which provides a number of advantages. First, Go supports cross-compilation for 32 and 64-bit systems for Mac, Linux and Windows. Go’s performance means that *vcfanno* can run large data sets relatively quickly. Go also offers a simple concurrency model, allowing *vcfanno* to perform intersections in parallel while minimizing the possibility of race conditions and load balancing problems that often plague parallel implementations. Moreover, as we demonstrate below, *vcfanno’s* parallel implementation of the chrom-sweep algorithm affords speed and scalability. Lastly, it is a very flexible tool because of its support for annotations provided in many common formats such as BED, VCF, GFF, BAM, and GTF.

## RESULTS

### Scalability of VCF annotation

We annotated the publicly available VCF files from both ExAC (v3; 10,195,872 VCF records) and the 1000 Genomes Project (Phase 3; 85,273,413 VCF records) to demonstrate *vcfanno’s* performance and scalability on both a whole-exome and a whole-genome dataset respectively. We used an extensive set of annotations and extracted a total of 50 different attributes from 17 distinct data sets representative of common annotations (see Supplementary File 1). We observed a near linear increase in annotation speed relative to a single core (3,302 and 7,457 seconds for ExAC and 1000 Genomes, respectively) when using 2-4 cores, but while performance continued to improve for additional cores, the improvement is sublinear (**Figure 4**). This is expected because we inevitably reach the limits of disk speed by concurrently accessing 17 files. Moreover, the degree of parallelism is limited by how fast the main process is able to read chunks of query VCF records that are kept in memory. Nonetheless, using 16 cores, *vcfanno* was able to annotate the variants from ExAC in less than 8 minutes and the 1000 Genomes variants in less than 17 minutes, performing at a rate of 21,902 and 85,452 variants per second, respectively.

**Figure 4.**
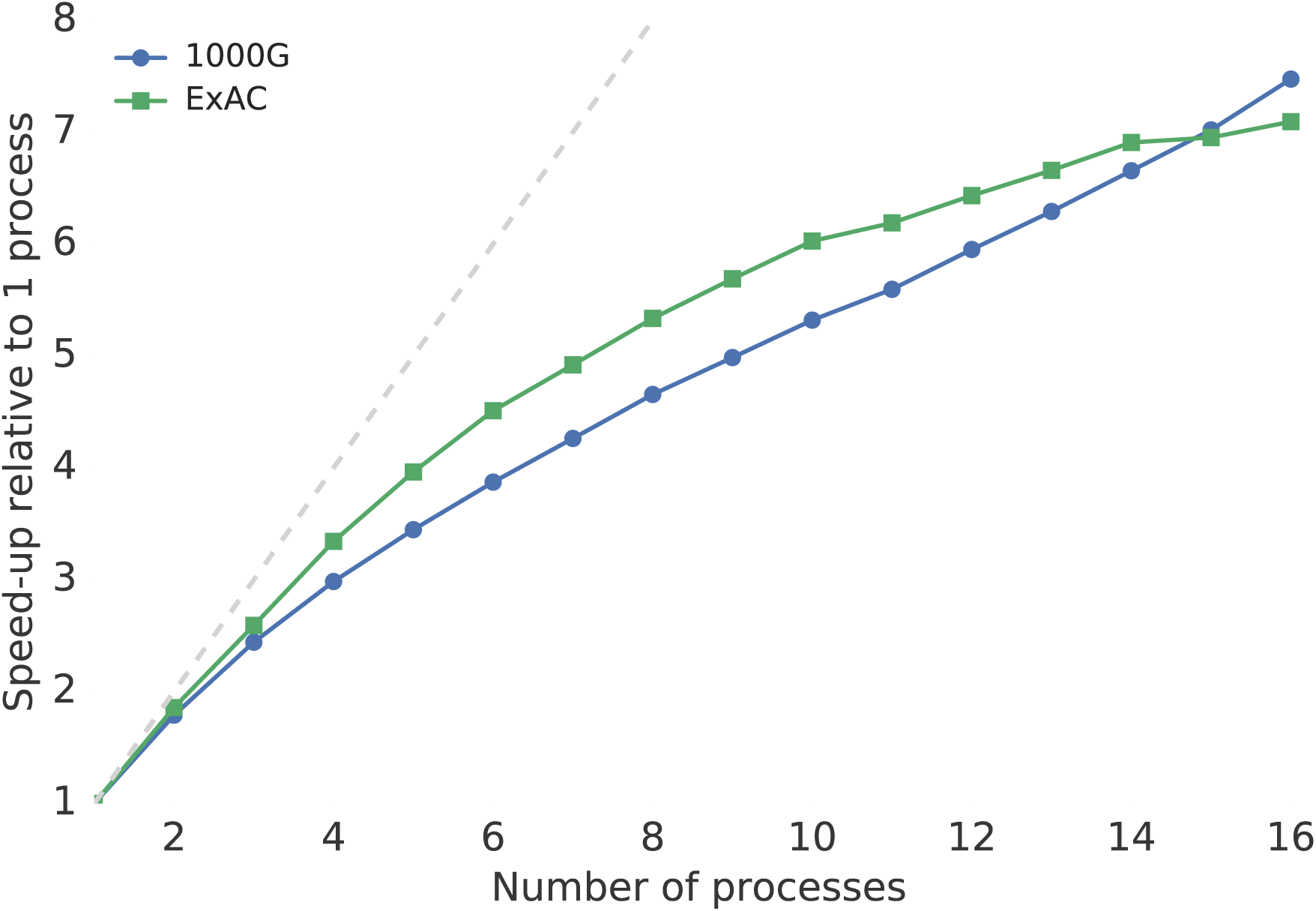
Parallelization efficiency. We show the efficiency of the parallelization strategy relative to 1 process on a whole-genome (1000 Genomes (1000G) in blue) and exome (ExAC in green) dataset. In both cases, we are short of the ideal speedup (gray line) but we observe a ~7-fold speedup using 16 processors. Absolute times are provided in Supplemental Table 1.

### The impact of interval distribution on performance

As described above, chunks of intervals constitute the individual units of work in the parallel “chrom-sweep” algorithm. A chunk is “full” when either the number of intervals in the array reaches the chunk size, or the genomic distance between two adjacent intervals is larger than the gap size. To understand the effect these parameters had on runtime, we varied both chunk size and gap size for the annotation of a whole-genome data set (variants from chromosome 20 of 1000 Genomes) and a whole-exome data set (variants from chromosome 20 of ExAC) given the same set of 17 annotation tracks from both whole-genome and whole-exome data sets that was used to create **Figure 4**. The choice of whole-genome and whole-exome query sets not only represents two common annotation use cases, but it also serves to illuminate the effect these parameters have on different data distributions. Not surprisingly, the run times for 1000 Genomes were completely dependent on chunk size and effectively independent of gap size, while ExAC runtimes exhibited the opposite behavior (**Figure 5A**). When intervals were more uniformly distributed throughout the genome, as with the 1000 Genomes data (**Figure 5B**), the distance between intervals tended to be small and therefore the maximum chunk size (not gap size) determined the number of intervals contained in the typical processing chunk. In contrast, since the intervals from ExAC tend to reside in smaller and more discrete clusters, maximum chunk size had almost no effect on the size of the typical processing chunk. As a result of this exploration, we have set the default chunk and gap sizes to work well on both whole-genome and exome datasets but we also allow them to be set by the user to maximize performance based on their knowledge of the data sets in question.

**Figure 5.**
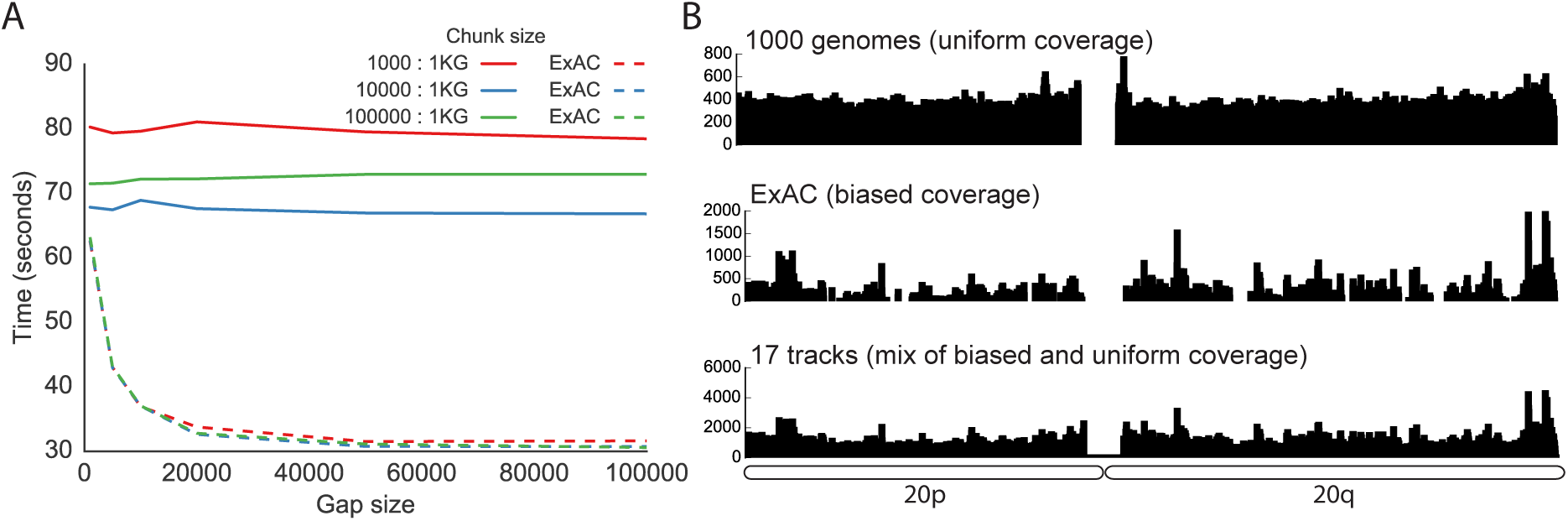
Effect of gap and chunk size on runtime for different data distributions. (A) Run-times for annotating small variants on chromosome 20 for 1000 Genomes (1KG) (1,822,268 variants) and ExAC (256,057 variants) against 17 annotation files using 4 cores and different combinations of gap size and chunk size. (B) Data density for chromosome 20 of 1000 Genomes, ExAC, and the summation of the 17 annotation files.

### Comparison to other methods

While no existing tools have the same functionality as *vcfanno*, BCFTools[18] includes an *annotate* command that allows one to extract fields from a *single* annotation file. Similarly, our own BEDTools[10, 11] uses the chrom-sweep algorithm to facilitate single-threaded intersection across multiple annotation files, yet it does not allow one to store annotations in the INFO field of the query VCF. Nonetheless, these tools provide an informative means to assess the performance of *vcfanno.* Using 9 different annotation sources ranging from whole-genome VCF to sparse BED files (see the *vcfanno* repository for replication code) we compared the runtime of *vcfanno* with 1, 4, 8, and 12 processes to that of BCFTools and BEDTools, both of which are single-threaded. We annotated the ExAC VCF with each tool. BEDTools can only intersect, not annotate, so we report the time to complete the intersections. BCFTools can only annotate 1 file at a time, so each of the nine annotations were conducted serially, and we report the total time required. BEDTools is an extremely efficient method for detecting interval intersections among multiple annotation files, but it is limited to a single core. BCFTools, on the other hand, can update the INFO field of the query with corresponding records from a single annotation file; but, it takes longer than *vcfanno*, even with a single process. Using 4 processors, *vcfanno* is 3.0 times as fast as BEDTools and 6.8 times as fast as BCFTools. The performance increase resulting from using eight processors is substantial, reducing the run-time from 703 seconds down to 591, but increasing to twelve processors yields little additional benefit.

**Table 1.** Speed comparison to other methods. We compare *vcfanno’s* performance to BEDTools and BCFTools using 1, 4, 8, and 12 processors when annotating the ExAC dataset using 9 annotation files. For all tools, we stream the output to bgzip in an effort to make the comparison as fair as possible. BCFTools can annotate only a single file at a time so the time reported is the sum of annotating each file and sending the result to the next annotation. This cannot be piped because the input to the BCFTools “annotate” tool must first be compressed by bgzip and subsequently indexed by Tabix.

| Method | Time (seconds) | No. Processors |
| --- | --- | --- |
| BEDTools | 2135.67 | 1 |
| BCFTools | 4776.55 | 1 |
| vcfanno | 2621.80 | 1 |
| vcfanno | 702.72 | 4 |
| vcfanno | 590.71 | 8 |
| vcfanno | 571.65 | 12 |

### Additional features

*Vcfanno* includes additional features that provide unique functionality with respect to existing tools. Annotating structural variants (SV) is complicated by the fact that, owing to the alignment signals used for SV discovery, there is often uncertainty regarding the precise location of SV breakpoints[19]. *Vcfanno* accounts for this uncertainty by taking into account the confidence intervals (defined by the CIPOS and CIEND attributes in the VCF specification) associated with SV breakpoints when considering annotation intersections. The confidence intervals define a genomic range in which the breakpoints are most likely to exist, therefore it is crucial for *vcfanno* to take these intervals into consideration when it considers annotations associated with SV breakpoints. Moreover, since SVs frequently affect hundreds to thousands of nucleotides, they will often intersect multiple intervals per annotation file. In such cases, the summary operations described above can be used to distill the multiple annotation intersections into a single descriptive measure.

Users will frequently need to further customize the annotations in the resulting VCF file. In order to facilitate this, *vcfanno* supports a concept of “post annotation”: that is, summary operations that are subsequently applied to the attributes that are extracted from annotation files for a given query VCF record. As an example, consider a situation where one would like to annotate each variant in one’s VCF file with the alternate allele frequency observed in the Exome Aggregation Consortium VCF. However, the ExAC VCF file solely provides the total count of chromosomes observed and the count of chromosomes exhibiting the alternate allele. Therefore, one cannot simply extract an alternate allele frequency directly from the ExAC VCF file. However, as illustrated in **Figure 6**, if the total and alternate allele counts are extracted with *vcfanno* (as “exac_total” and “exac_alts” below), one can define an additional “post-annotation” section that uses lua to compute the alternate allele frequency (“exac_aaf”) from the total and alternate allele counts extracted from the ExAC VCF file. Users can write extensive lua functions in an external script and subsequently call these in the *annotation* and *post-annotation* sections of the configuration file. For example, in **Figure 6C**, we use an external lua script (provided to *vcfanno* on the command line) to implement a function (see Supplementary File 2) that calculates the lower bound of the allele frequency confidence interval. This is useful for determining whether an allele frequency is different from 0 based on the 95% confidence bounds. This type of specialized logic is simple to implement given lua’s scripting capabilities and allows *vcfanno* to be customized to a researcher’s specific needs. In fact, this “post-annotation” concept can be applied “in place” to a VCF without any annotation files, thereby allowing the user to perform modifications to a VCF file’s INFO field.

**Figure 6.**
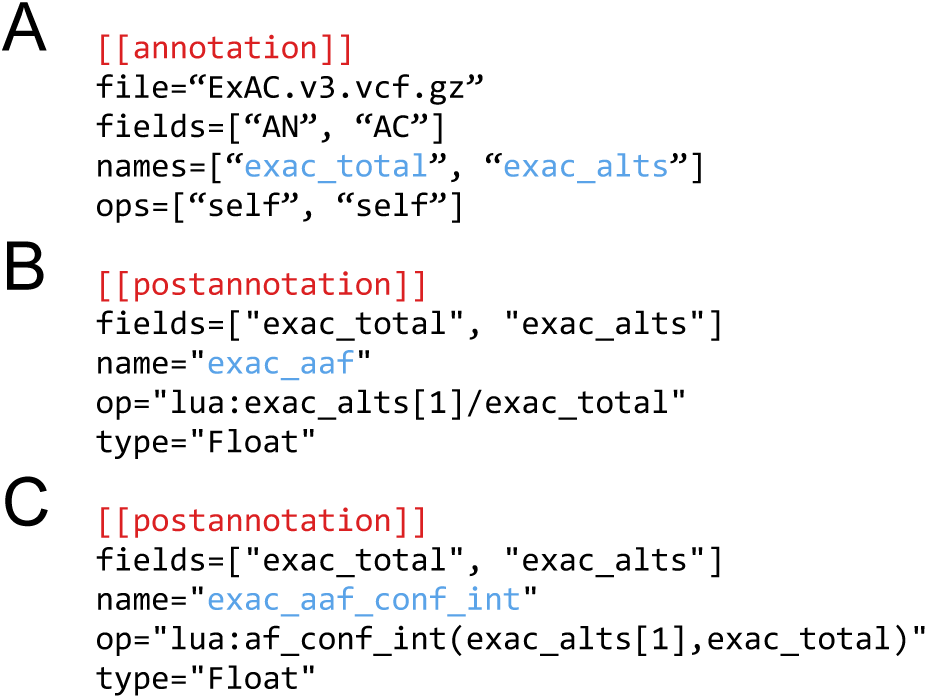
Using a “post-annotation” block to compute new annotations derived from existing annotations. **(A)** As described in the text, computing this post-annotation assumes that the “exac_total” and “exac_alts” fields have been extracted (via the AN and AC fields in the ExAC VCF) from the annotation file using standard annotation blocks. The AC field returns an array of alternate alleles for the variants in the annotation file that match a given variant in the query VCF. **(B)** In this post-annotation block, we calculate the alternate allele frequency in ExAC (“exac_aaf”) using “exac_alts[1]” as the numerator because in this example we calculate the alternate allele frequency based on the first alternate allele. **(C)** An example of a post-annotation block that calls a function (af_conf_int) in an external lua script (see Supplemental File 2) to compute the lower bound of the allele frequency confidence interval based upon counts of the alternate allele and total observed alleles. (D) An example invocation of vcfanno that annotates a BGZIP-compressed VCF file (example.vcf.gz) using the configuration file described in panels A-C (red), together with the lua script file (blue) containing the code underlying the af_conf_int function.

## DISCUSSION

We have introduced *vcfanno* as a fast and flexible new software resource that facilitates the annotation of genetic variation in any species. We anticipate that *vcfanno* will be broadly useful to researchers both as a standalone annotation tool, and also in conjunction with downstream VCF filtering and manipulation software such as snpEff[6], BCFTools[18], BGT[20], and GQT[21]. There are, however, caveats to the proper use of *vcfanno*, and it exhibits poorer performance in certain scenarios. First, when annotating with other VCF files, it is recommended that both the variants in the query VCF and each database VCF are normalized and decomposed in order to ensure that both variant sites and alleles are properly matched when extracting attributes from the database VCF files[22]. Secondly, *vcfanno’s* relative performance is, not surprisingly, less impressive on very sparse datasets (e.g., 1 or 2 variants every 20KB) such as a VCF resulting from the exome sequencing of one individual. While annotating these files is still quite fast, typically between three and five minutes, the sparsity of data exposes an exposes the overhead associated with using Tabix to create streams of database intervals that are germane to the current chunk. Tabix must decompress an entire BGZF block from each annotation file even if the query chunk merely includes a single variant because Tabix’s smallest BGZF block represents a genomic range of 16 kilobases. Therefore, when the query VCF is very sparse, an entire BGZF block from each annotation is frequently (and wastefully) decompressed for each query variant. In future versions of *vcfanno*, we will explore alternative approaches in order to avoid this limitation, thereby maximizing performance in all usage scenarios.

## CONCLUSION

*Vcfanno* is an extremely efficient and flexible software package for annotating genetic variants in VCF format in any species. It represents a substantial improvement over existing methods, enabling rapid annotation of whole-genome and whole-exome datasets and provides substantial analytical power to studies of disease, population genetics, and evolution.

## ACKNOWLEDGMENTS

We acknowledge Liron Ganel for helpful suggestions in developing support for annotating structural variants.

## FUNDING

This research was supported by a US National Human Genome Research Institute award to A.R.Q. (NIH R01HG006693).

## AUTHOR CONTRIBUTIONS

B.S.P. implemented the software, analyzed the data, and wrote the manuscript. R.M.L. contributed to the design of the parallel algorithm, analyzed the data, and wrote the manuscript. A.R.Q. conceived of the software and wrote the manuscript.

## COMPETING INTERESTS

The authors declare no competing interests.

## REFERENCES

1. Danecek P, Auton A, Abecasis G, Albers CA, Banks E, DePristo MA, Handsaker RE, Lunter G, Marth GT, Sherry ST, McVean G, Durbin R, 1000 Genomes Project Analysis Group: The variant call format and VCFtools. Bioinformatics 2011, 27:2156–2158.

2. McKenna A, Hanna M, Banks E, Sivachenko A, Cibulskis K, Kernytsky A, Garimella K, Altshuler D, Gabriel S, Daly M, DePristo MA: The Genome Analysis Toolkit: a MapReduce framework for analyzing next-generation DNA sequencing data. Genome Res 2010, 20:1297–1303.

3. Garrison E, Marth G: Haplotype-based variant detection from short-read sequencing. arXiv [q-bio.GN] 2012.

4. Li H: Toward better understanding of artifacts in variant calling from high-coverage samples. Bioinformatics 2014, 30:2843–2851.

5. Koren A, Polak P, Nemesh J, Michaelson JJ, Sebat J, Sunyaev SR, McCarroll SA: Differential relationship of DNA replication timing to different forms of human mutation and variation. Am J Hum Genet 2012, 91:1033–1040.

6. Cingolani P, Platts A, Wang LL, Coon M, Nguyen T, Wang L, Land SJ, Lu X, Ruden DM: A program for annotating and predicting the effects of single nucleotide polymorphisms, SnpEff: SNPs in the genome of Drosophila melanogaster strain w1118; iso-2; iso-3. Fly 2012, 6:80–92.

7. Wang K, Li M, Hakonarson H: ANNOVAR: functional annotation of genetic variants from high-throughput sequencing data. Nucleic Acids Res 2010, 38:e164.

8. McLaren W, Pritchard B, Rios D, Chen Y, Flicek P, Cunningham F: Deriving the consequences of genomic variants with the Ensembl API and SNP Effect Predictor. Bioinformatics 2010, 26:2069–2070.

9. Exome Aggregation Consortium, Lek M, Karczewski K, Minikel E, Samocha K, Banks E, Fennell T, O’Donnell-Luria A, Ware J, Hill A, Cummings B, Tukiainen T, Birnbaum D, Kosmicki J, Duncan L, Estrada K, Zhao F, Zou J, Pierce-Hoffman E, Cooper D, DePristo M, Do R, Flannick J, Fromer M, Gauthier L, Goldstein J, Gupta N, Howrigan D, Kiezun A, Kurki M, et al.: Analysis of protein-coding genetic variation in 60,706 humans. bioRxiv 2015:030338.

10. Quinlan AR, Hall IM: BEDTools: a flexible suite of utilities for comparing genomic features. Bioinformatics 2010, 26:841–842.

11. Quinlan AR: BEDTools: The Swiss-Army Tool for Genome Feature Analysis. Curr Protoc Bioinformatics 2014, 47:11.12.1–11.12.34.

12. Neph S, Kuehn MS, Reynolds AP, Haugen E, Thurman RE, Johnson AK, Rynes E, Maurano MT, Vierstra J, Thomas S, Sandstrom R, Humbert R, Stamatoyannopoulos JA: BEDOPS: high-performance genomic feature operations. Bioinformatics 2012, 28:1919–1920.

13. Kent WJ, Sugnet CW, Furey TS, Roskin KM, Pringle TH, Zahler AM, Haussler D: The human genome browser at UCSC. Genome Res 2002, 12:996–1006.

14. Layer RM, Quinlan AR: A Parallel Algorithm for $N$-Way Interval Set Intersection. Proc IEEE:1–10.

15. McKenney M, McGuire T: A parallel plane sweep algorithm for multi-core systems. In Proceedings of the 17th ACM SIGSPATIAL International Conference on Advances in Geographic Information Systems. ACM; 2009:392–395.

16. Khlopotine AB, Jandhyala V, Kirkpatrick D: A Variant of Parallel Plane Sweep Algorithm for Multicore Systems. IEEE Trans Comput Aided Des Integr Circuits Syst 2013, 32:966–970.

17. Daniel Kortschak R, Adelson DL: bíogo: a simple high-performance bioinformatics toolkit for the Go language. bioRxiv 2014:005033.

18. Li H, Handsaker B, Wysoker A, Fennell T, Ruan J, Homer N, Marth G, Abecasis G, Durbin R, 1000 Genome Project Data Processing Subgroup: The Sequence Alignment/Map format and SAMtools. Bioinformatics 2009, 25:2078–2079.

19. Quinlan AR, Hall IM: Characterizing complex structural variation in germline and somatic genomes. Trends Genet 2012, 28:43–53.

20. Li H: BGT: efficient and flexible genotype query across many samples. Bioinformatics 2016, 32:590–592.

21. Layer RM, Kindlon N, Karczewski KJ, Exome Aggregation Consortium, Quinlan AR: Efficient genotype compression and analysis of large genetic-variation data sets. Nat Methods 2015.

22. Tan A, Abecasis GR, Kang HM: Unified representation of genetic variants. Bioinformatics 2015, 31:2202–2204.

23. Li H: Tabix: fast retrieval of sequence features from generic TAB-delimited files. Bioinformatics 2011, 27:718–719.

